# Flight power muscles have a coordinated, causal role in hawkmoth pitch turns

**DOI:** 10.1101/2023.09.27.559785

**Authors:** Leo Wood, Joy Putney, Simon Sponberg

**Author notes:** **Corresponding Author:** Simon Sponberg.

## Abstract

Flying insects solve a daunting control problem of generating a patterned and precise motor program to stay airborne and generate agile maneuvers. In this motor program consisting of every action potential controlling wing musculature, each muscle encodes significant information about movement in precise spike timing down to the millisecond scale. While individual muscles share information about movement, we do not yet know if they have separable effects on an animal’s motion, or if muscles functionally interact such that the effects of any muscle’s timing depends heavily on the state of the entire musculature. To answer these questions, we performed spike-resolution electromyography and precise stimulation of individual spikes in the hawkmoth *Manduca sexta* during tethered flapping. We specifically explored how the flight power muscles themselves may contribute to pitch control which is necessary to stabilize flight. Combining correlational study of visually-induced turns with causal manipulation of spike timing, we discovered likely coordination patterns for pitch turns, investigated if these correlational patterns can individually drive pitch control, and studied whether the precise spike timing of indirect power muscles can lead to pitch maneuvers. We observed significant timing change of the main downstroke muscles, the dorsolongitudinal muscles (DLMs), associated with whether a moth was pitching up or down. Causally inducing this timing change in the DLMs with electrical stimulation produced a consistent, mechanically relevant feature in pitch torque, establishing that indirect power muscles in *Manduca* have a control role in pitch. Because changes were evoked in unconstrained flapping in only the DLMs, however, these pitch torque features left large unexplained variation. We find this unexplained variation indicates significant functional overlap in pitch control such that precise timing of one power muscle does not produce a precise turn, demonstrating the importance of coordination across the entire motor program for flight.

**Summary Statement:** We investigate how individual muscles contribute to flight by manipulating muscle timing in behaving hawkmoths. We find precise timing of single muscles does not produce precise turns, highlighting the importance of coordination across the entire motor program.

## INTRODUCTION

Locomoting animals have to generate a motor program, a coordinated spatial and temporal pattern of activity sent to an array of muscles to produce behaviors. Each muscle is controlled by action potentials, or spikes, which across many animals carry information about the motion of an animal to a highly precise millisecond or sub-millisecond scale (Sober et al., 2018; Putney et al., 2019). This is significant because the degree to which muscles are *functionally* coordinated, such that the kinematic and behavioral action of individual muscles depends on the action of other muscles, determines how precise spike timing translates to movement. If a single muscle independently controls specific features of kinematics or dynamics, then precisely timed spikes to that muscle can be directly attributed to precise behavioral outcomes, as happens in *Drosophila* with the control of pitch by the basalar muscles (Whitehead et al., 2022). But if control is orchestrated across many muscles simultaneously such that the action of one muscle changes the potential of another muscle to do control, then the transformation of a precise change in spike timing into movement will depend on the context created by the spiking patterns in the rest of the motor program. Such a dependency might be expected since muscles often act in coordinated groups

For one flying insect species, the hawkmoth *Manduca sexta*, the flight motor program is already known to be sub-millisecond precise (Putney et al., 2019; 2021b), and correlational evidence suggests a high degree of coordination and overlapping control potentials. The hawkmoth motor program is information-theoretic coordinated so that spike timing of all muscles carries redundant global information about behavioral output(Putney et al., 2019). Linear combinations of muscle activity also explain the behavioral output of multiple different turning maneuvers, allowing decoding with 90% accuracy the type of turn maneuver performed from activity of only 4 or more muscles (Putney et al., 2021a). These findings stem from hawkmoths providing a particularly tractable system for understanding motor precision and coordination. Their flight musculature has comparatively few muscles, all effectively single motor units (Rheuben, 1985; Usherwood, 1962), so a comprehensive motor program can be obtained through electrophysiology on a behaving animal from the activity of only 10 muscles (Putney et al., 2019). Like many flying insects, this motor program is separated into indirect flight muscles and a set of steering muscles that directly attach to the wing, all contained within the thorax of the body. Two large pairs of flight power muscles, the dorsolongitudinal muscles (DLMs) and dorsoventral muscles (DVMs), produce most of the mechanical power for flight, with small steering muscles attached directly to the wing hinge adjusting wing kinematics (Eaton et al., 1988). Steering muscles are well known to perform flight control, with timing of individual muscles often correlated to specific maneuvers and wing kinematics (Ando and Kanzaki, 2004; Wang et al., 2008; Hedenström, 2014; Dickinson and Tu, 1997; Tu and Dickinson, 1996; Balint and Dickinson, 2004; Sadaf et al., 2015; Lindsay et al., 2017). But with the timescale of power muscle force production well within a single wingstroke (Tu and Daniel, 2004), hawkmoths provide a case where power and steering muscles both can be actively involved in flight control (Sponberg and Daniel, 2012; Sponberg et al., 2015a; Ando and Kanzaki, 2016).

Hawkmoths, then, possess a precisely timed motor program (Putney et al., 2019; 2021b) using both power and steering muscles for control (Sponberg and Daniel, 2012; Sponberg et al., 2015a; Ando and Kanzaki, 2016), and there is correlational evidence suggestive of coordination and overlapping control potentials (Putney et al., 2019; 2021a). But when the spike timing of *many* muscles may be correlated with behavior, it is challenging to separate the causal contributions of individual muscles to a given behavioral output. Direct electrical stimulation of muscles and motor neurons can reveal a muscle’s control potential, by causally inducing specific spike timing in specific situations. As an *in vivo* manipulation of muscle activity, electrical stimulation has been used to identify muscle control potentials in anesthetized vertebrates (Vazquez, 1995), elicit control changes in flying insects via high frequency stimulation (Sato et al., 2015; Sato and Maharbiz, 2010), or perform bulk perturbation to specific sides of an insect (Tsang et al., 2010). More rarely, however, it can be applied in behaving invertebrates to produce targeted, spike-level manipulations of specific muscle timings (Sponberg et al., 2011). In hawkmoths, altering motoneuron spike timing shows a causal connection between left-right timing differences in primary downstroke muscles and yaw torque (Sponberg and Daniel, 2012). However, for flying insects these temporally precise manipulations and related correlational studies have only focused on left-right asymmetries leading to roll or yaw turns (Springthorpe et al., 2012; **?**; Wang et al., 2008; Sponberg and Daniel, 2012; Sponberg et al., 2015a). Causal manipulation of steering muscles to produce pitch turns has been performed in fruit flies using optogenetics (Whitehead et al., 2022), but for flying insects no spike-level manipulations have been performed to study pitch turns.

Gaps such as these are notable because pitch turns are particularly interesting for hovering insects. Nearly every model of insect flight dynamics identifies an unstable mode resulting from coupled oscillations in fore-aft velocity and pitch, requiring some degree of active neuromuscular control (Windsor et al., 2014; Ristroph et al., 2013; Kim and Han, 2014; Kim et al., 2015). Active control of pitch must also be bilaterally symmetric by nature, so any control cannot simply result from left-right motor program asymmetries (Jankauski et al., 2017). Though flying insects such as *Dipterans* and *Lepidopterans* control pitch through a multifaceted set of passive and active mechanisms, including passive vibrational stability via wing oscillation (Taha et al., 2020) and movement of the center of mass (COM) relative to the center of pressure (COP) via abdominal motion (Ristroph et al., 2013; Frye, 2001; Hinterwirth and Daniel, 2010; Hedrick and Daniel, 2006), most pitch control is thought to arise from the activity of the steering muscles (Whitehead et al., 2015; 2022). While modulation of spike timing in the flight power muscles has been closely linked to roll and yaw turns (Sponberg and Daniel, 2012; Sponberg et al., 2015a; Springthorpe et al., 2012; Lehmann et al., 2013), it is not known to what degree their potential to control extends into pitch turns. Such turns require not left-right phase separation but modulation of kinematics between the downstroke and upstroke. If the indirect power muscles are capable of contributing to pitch control, they must do so through alteration of wing kinematics, either moving the COP, changing time-varying wing forces, or producing inertial moments on the COM. In freely flying hawkmoths pitching up in response to looming stimuli correlates with changes in angle of attack, stroke plane tilt, angle at wingstroke reversals, and deviation angle at wingstroke reversals (Cheng et al., 2011). The interval between the DLM and DVM has been implicated in changing deviation angle of the wing in free flight (Wang et al., 2008), and shown to change with flight speed in forward flight, so may be related to modulating power production (Hedrick et al., 2017).

Overall, however, while the precise spike timing of the hawk-moth power muscles has been connected to asymmetric kinematics, their potential for control on other maneuvers such as pitch turns are unclear. We hypothesize that the indirect power musculature in the hawkmoth *Manduca sexta* does contribute to pitch control. We also hypothesize that the flight motor program is functionally coordinated, such that even the mapping of individaul action potentials of large power muscles into movement (the muscle’s control potential) (Sponberg et al., 2011) is dependent on the context of the rest of the motor program. Alternatively, we might see that the flight motor program is not very functionally coordinated, so that the control potentials of individual muscles are easily identified and not affected by the activity of other muscles. Using a combination of correlational and causal experiments, we test both of these hypotheses, identifying an association between power muscle activity and pitch turns, then rewriting motor activity to reproduce those patterns in tethered flight.

## MATERIALS AND METHODS

### Animals

All moths (*M. sexta*) were obtained as pupae and housed communally after eclosion with a 12 hour light-dark cycle (N = 9, University of Washington colony and Carolina Biological Supply Co for visually-induced turning experiments; N = 5, Case Western Reserve colony for stimulation experiments). Naive males and females were used in experiments conducted during the dark period of their cycle. EMG recordings from flight muscles were performed following the same procedure as (Putney et al., 2019), where moths were cold-anesthetized and pairs of silver wire (diameter 0.005*in*) were inserted into the thorax and fastened with cyanoacrylate glue. After wire insertion, moths were fastened with cyanoacrylate to a tether rigidly attached to a custom six-axis load cell (ATI Nano17ti, FT20157; calibrated ranges *F*_*x*_, *F*_*y*_ =*±* 1.0*N, F*_*z*_ =*±* 1.8*N, τ*_*x*_, *τ*_*y*_, *τ*_*z*_ =*±* 6.25 *mNm*). After attachment moths were left to dark adapt for 30 minutes at luminance levels typical for when these crepuscular moths are active (Sponberg et al., 2015b).

### Hard turns in visual arena experiments

The data for the correlational visually-induced hard turns experiment of this study were originally published by Putney et al (Putney et al., 2021a; **?**). Moths were presented with wide-field sinusoidal gratings on a rendered 3D sphere in Microsoft Visual Studio (Fig. 1). The stimuli were projected onto three computer monitors (ASUS PG279Q ROG Swift; 2560 × 1440 px) overlaid with neutral density filters to obtain peak sensitivity luminance conditions for *Manduca* of approximately 1 *cd/m*^2^ (Stöckl et al., 2017). The tethering set-up placed the moth at the center of a three-sided box formed by these three monitors (Fig. 1A). The sinusoidal gratings had a spatial frequency of 20°/cycle, and the sphere was rotated to place its axis of rotation along each of the earth-coordinate system flight axes of pitch, roll, and yaw (Fig. 1B). The sphere was then rotated about this axis in opposite directions at a constant drift velocity of 100 °/s (5 cycle/s), which was chosen since this spatiotemporal frequency drove high responses in the moth visual system (Stöckl et al., 2017). Therefore, each moth responded to six distinct visual stimulus conditions, three pairs about each flight axis: pitch up (PU), pitch down (PD), roll left (RL), roll right (RR), yaw left (YL), and yaw right (YR).

**Fig. 1.**
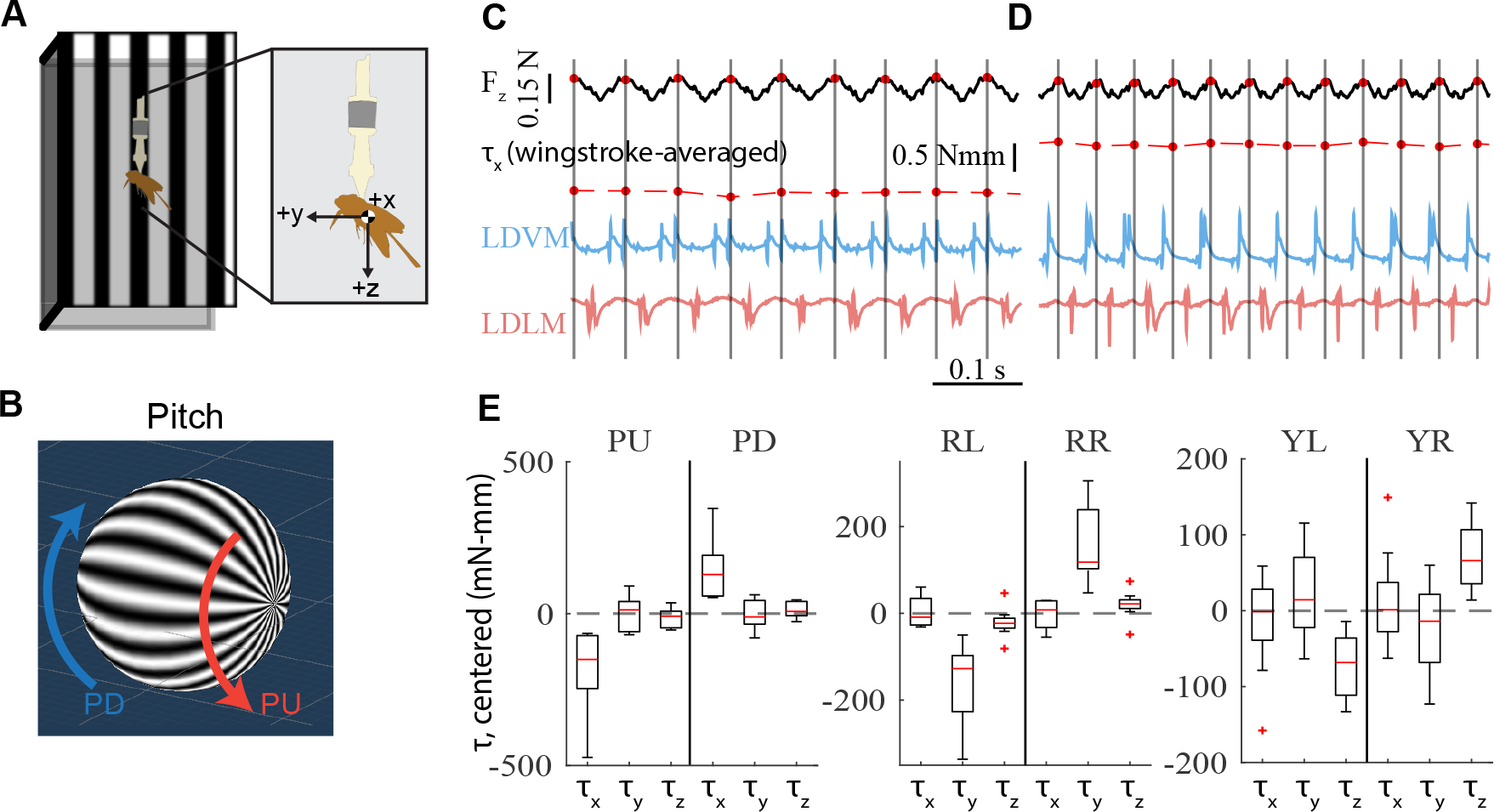
Visually induced hard turns produce separated body torque responses. Tethered moths were shown three pairs of visual stimuli to elicit hard turns about the three flight axes – pitch, roll and yaw. (**A**) Schematic of the visual stimulus arena formed by three computer monitors, with the moth tethered in the center. Inset depicts coordinate frame used throughout. (**B**) Example of wide-field stimuli in pitch direction. Wide-field sinusoidal gratings on a projected 3D sphere, with spheres rotated with a constant drift velocity produce two different stimulus conditions for each axis of rotation. (**C-D**) Simultaneous raw EMG recordings of the left side power muscles with motor output for 0.5 s in the pitch up (C) and pitch down (D) conditions. Wingstrokes were segmented using the Hilbert transform of *F*_*z*_ force. Wingstroke averaged pitch torque (*τ*_*x*_) demonstrates different behavior in response to the visual stimulus conditions. (**E**) Centered mean wingstroke averaged pitch (*τ*_*x*_), roll (*τ*_*y*_), and yaw (*τ*_*z*_) torques for each individual moth (N = 9) in six conditions.

Simultaneous strain gauge voltage recordings from the F/T transducer and EMG recordings from the silver wire electrodes were taken as the moth responded to each of the six stimulus conditions for 20 seconds, each recorded at 10000 Hz. The strain gauge voltages were converted to the three forces and three torques on each axis at the estimated average COM for tethered moths. These forces and torques were lowpass filtered using a 8th order Butterworth filter with a cutoff of 1000 Hz. To segment wing strokes in some analyses, a type II Chebychev filter was applied to *F*_*z*_ with a bandpass between 5 and 35 Hz to capture the range of wingbeat frequencies observed in this dataset (15 - 27 Hz). The Hilbert transform of this filtered signal was taken to estimate the wingstroke phase, and individual wingstrokes were separated by finding negative-to-positive zero crossings of this phase, with t=0 for each wingstroke occurring at these crossings. This use of the Hilbert transform to find instantaneous phase and separate a gait into specific cycles has previously been used in hawkmoths (Sponberg et al., 2015a; Putney et al., 2019), gait analysis of cockroaches (Revzen and Guckenheimer, 2008), and rat whisking (Hill et al., 2011). Spikes in the raw EMG voltage recordings were discriminated using Offline Sorter (OFS; Plexon) via threshold crossing. Where necessary, filtering options in this software were used to correct baseline wander, motion artifacts, and other noise that made discriminating spikes consistently difficult. Spike times were specified to 0.1 ms relative to zero phase in each wing stroke as described above.

Centered wingstroke mean torques were used to compare between opposing conditions in pitch, roll, and yaw (Fig. 1E). Centering was performed by subtracting from per-wingstroke mean pitch, roll, and yaw torques the per-moth overall mean pitch, roll, and yaw torques *τ*_*x*_, *τ*_*y*_, *τ*_*z*_ across all that moth’s data for both conditions (i.e. both pitch up and pitch down turns). Centering was performed independently for each moth (*N* = 9 moths).

### Electrical stimulation experiments

To causally manipulate power muscle timing and observe resulting changes in flight forces and torques, electrical stimulation was used to alter the spike timing of the dorsolongitudinal muscles (DLMs). While there are many previous studies eliciting control changes in flying insects using high frequency electrical stimulation (Sato et al., 2015; Sato and Maharbiz, 2010) or bulk stimulation of individual sides of an insect (Tsang et al., 2010), here we use targeted and precisely timed stimulation to induce single action potentials in the DLMs, in a method more akin to (Sponberg et al., 2011) and (Sponberg and Daniel, 2012).

Moths attached to a 6-axis load cell tether would freely engage in bouts of flapping flight, during which the DLMs would be stimulated with a specific timing relative to the DVMs. Stimulus delay times were chosen at random from a range of 4-40 ms after DVM spike detection, with each delay time applied for a 20 second recording period with EMG and F/T transducer recorded at 10 kHz. Single 0.25 ms pulses of either constant current or constant voltage were applied to silver wires placed on either end of each DLM, with each stimulation separated by at least 4 seconds to avoid entrainment. A diagram signal flow used for stimulation is shown in Fig. 2A. In brief, timing of stimulus was achieved by passing either the left or right DVM voltage recording through a custom analog spike-detection circuit and microcontroller. The controller would trigger a stimulator (A-M Systems Model 3800) with a stimulus isolation unit (A-M Systems Model 2200) on a controlled delay time from the onset of a detected DVM spike.

**Fig. 2.**
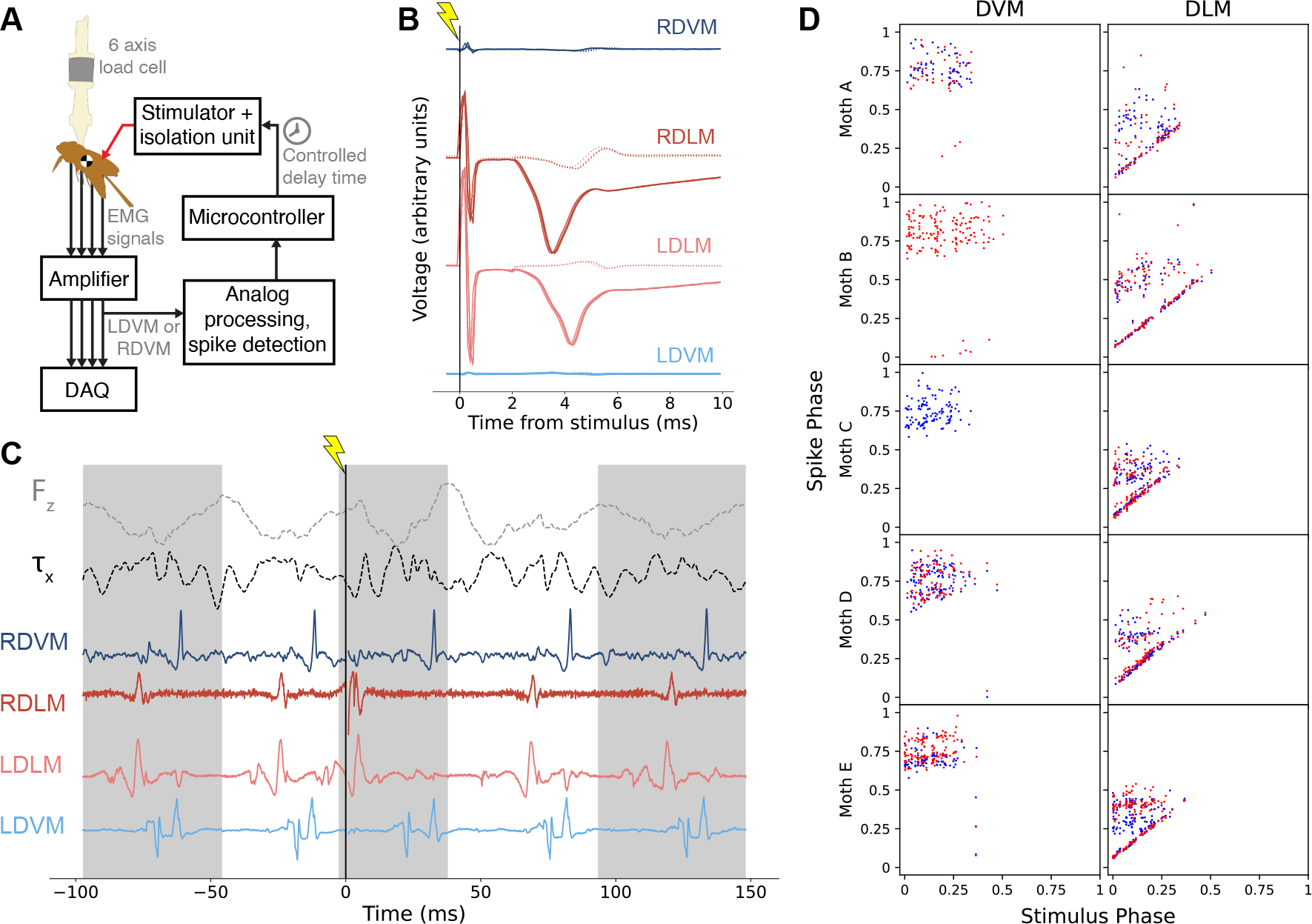
Electrical stimulation allows controlled change of DLM timing, leading to changes in body pitch torque. (**A**) Diagram of stimulation experiment signal flow. Analog spike detection is run on a DVM signal post-amplification, enabling a microcontroller to trigger a stimulator with specific timings relative to observed DVM spikes (see (Sponberg and Daniel, 2012) for a similar experiment). (**B**) Voltage recordings, aligned at time of stimulus, from all four power muscles in a quiescent moth while stimulation is applied, illustrating efficacy of electrical stimulation. Lighter dotted traces are 0.03mA current of stimulation, darker solid traces are 0.05mA current. (**C**) Example trace of EMG and vertical force and pitching torque data during stimulation. Stimulation is applied at time *t* = 0 (denoted by vertical line with lightning symbol) and produces a phase advance in RDLM and LDLM spikes. Alternating shaded regions indicate wingstrokes found via negative-to-positive zero crossings of the Hilbert transform of *F*_*z*_ (light grey dashed line). (**D**) Phase of all spikes in stimulation wingstroke plotted against the phase at which stimulation was applied for that wingstroke on the x axis. Red points indicate right muscle, blue points left muscle. If stimulation is inducing spikes in the DLM, then spikes along an identity line should be observed only in the right DLM column.

All stimulus pulses were biphasic, but specific stimulus features such as whether constant current or voltage were used and of what amplitude, were calibrated for each moth individually. This calibration in an example moth is shown in Fig. 2B. Stimulus was repeatedly applied while the moth was in a quiescent state, with stimulation amplitude steadily increased until evoked motor action potentials were observed in both DLMs with no evoked action potentials observed in surrounding muscles (Fig. 2B). While most individuals had their best response from constant current stimulation, differences in electrode placement and individual anatomy and physiology resulted in better results for some individuals from constant voltage stimulation. After calibration, stimuli ranged from 0.05-0.1 mA for constant current stimulation and 5-10 V for constant voltage. Note that, as shown in Fig. 2B, evoked action potentials occur roughly 4 milliseconds after a pulse of current is applied.

Spike discrimination, wingstroke segmentation, and FT data calibration and filtering followed the same parameters as described previously (ADD CITE), with several exceptions. The moth’s tether is not located precisely at the center-of-mass (COM). Strain gauge voltages were transformed to COM locations estimated for each individual by performing bounded linear least-squares minimization to find the COM location which minimized torques to zero during fully quiescent data of each tethered moth. Different filter parameters were used for wingstroke segmentation, specifically a 4th order type II Chebyshev filter with a 10-40 Hz passband, as this demonstrated better wingstroke segmentation performance for the stimulation data.

Stimulation trials were included in this study only if stimulus evoked a muscle action potential (MAP) in both DLMs, and if the first observed MAP from both DLMs in the stimulus wingstroke was evoked rather than natural. MAPs were deemed evoked rather than natural if they occurred within a 5ms window directly following stimulation. Figure 2D provides a visual aide for how evoked MAPs were selected; wingstrokes were only included if both left and right DLMs had a MAP occur along the main diagonal of Fig. 2D, and if the first spike in the stimulation wingstroke occurred along this diagonal.

### Feature extraction and stimulation analysis

To analyze the effects of controlled DLM timing on pitch torque, features of pitch torque which were associated with evoked DLM timing had to be extracted and quantified. To extract features of pitch torque that correlated with stimulation-evoked changes in DLM timing, canonical correlation analysis (CCA) was employed with the cross decomposition module of the *scikit-learn* python package (Pedregosa et al., 2011), which utilizes a previously described algorithm (Wegelin, 2000). CCA is one of several dual-dimensionality reduction methods, such as partial least squares regression (PLS) (Wegelin, 2000; Sponberg et al., 2015a), where two sets of latent variables (or “features”) ***τ*** ^*′*^ and ***Y*** ^***′***^ which maximally correlate with each other are extracted from two sets of original data variables ***τ*** and ***Y***. CCA is very closely related to the commonly used principal components analysis (PCA) in that it uses an eigendecomposition to construct components (or “features”) from a linear combination of variables in a dataset. In PCA, these features are constructed to maximize variance, whereas in CCA and the broader family of PLS, two sets of features which maximally correlate with each other are constructed simultaneously from two sets of variables. By inputting the set of pitch torque waveforms for ***τ*** and the stimulation-evoked DLM timing or phase for ***Y***, we can extract a feature of the pitch torque data, ***τ***, which maximally correlates with evoked timing changes.

CCA was performed on each individual moth independently with pitch torque data assembled into a matrix ***τ*** and stimulation-evoked DLM phase assembled into a matrix ***Y***. Pitch torques for all stimulation wingstrokes were linearly interpolated to *m* = 300 samples long and assembled into an *n×m* matrix ***τ***, where *n* is the number of valid stimulation wingstrokes for that individual. The other CCA input, ***Y***, was constructed as an *n×k* matrix of just a single variable, the phase of the evoked DLM spike for each stimulation wingstroke (*k* = 1 for the duration of this paper).

Note that CCA was performed on z-scored versions of ***τ***and ***Y***, denoted throughout by caret hats as 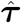 and ***Ŷ***, respectively. Z-scoring allows for more stable operation on data of different ranges and units by making variance non-dimensional and placing disparate data in similar ranges. Note that when quantities are provided in real units z-scoring is inverted.

For ***τ*** and ***Y***, CCA produces a *k×m* set of features often referred to as “loadings” *τ* _*loadings*_, an *m×k* set of “weights” used to construct the features *τ*_*weights*_, and an *n×k* set of “scores” defined by ***τ****τ* _*weights*_, effectively the projection of the data in ***τ*** to the reduced set of features ***τ***^*′*^. Data in ***τ*** can be reconstructed from the features in *τ*_*loadings*_

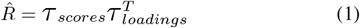

where 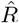 is an *n×m* matrix of approximations of the original data in ***τ*** via linear scaling of the feature vector *τ*_*loadings*_. Note that, denoted by the caret hat, 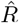 is still z-scored, and the inverse of the z-scoring is applied to obtain values with real units.

Angular impulse, or net change in angular momentum, was calculated from pitch torque of extracted features and original pitch torque data for individual wingstrokes as

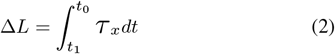

where *t*_0_ and *t*_1_ define the start and end times of the wingstroke, respectively. We obtain effective pitch angular velocity due to pitch torque as

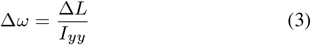

where *I*_*yy*_ is the pitch moment of inertia for *Manduca sexta*, defined throughout this work as *I*_*yy*_ = 266.7 *gmm*^2^ from (Cheng et al., 2011). Numerical integration was performed with the trapezoidal method, with CCA feature reconstructions resampled from *m* = 300 to the original number of samples present in the corresponding wingstroke.

## RESULTS

We investigated the relationship between precise spike timing of the flight power muscles in *Manduca sexta* and pitch turns through two main approaches. In the first experiment, *n* = 21,203 total wingstrokes were analysed from *N* = 9 moths induced to perform roll, pitch, and yaw turns through rotating wide-field visual stimuli to correlate how timing of the primary flight muscles changed between pitch up and pitch down turns. In the second experiment, in *n* = 345 wingstrokes across *N* = 5 moths we causally manipulated the timing of the primary downstroke muscles, the DLMs, to observe changes in pitch torque due to altering the relative timing of the primary flight power muscles.

### Power muscle spike timing correlates with pitch turns in visually-induced turns

Mean centered torques *τ* ^*′*^ for each condition showed separation on the expected axis (Fig. 1E), demonstrating that each condition indeed elicited a strong turn response from moths in only the desired direction. For instance, overall mean wingstroke torques per individual moths (*N* = 9) were statistically significantly different between mean pitch torque *τ* _*x*_ (*p* = 0.003 in paired t-test), but not between mean roll or yaw torques (*p >* 0.05 in paired t-tests for *τ* _*y*_ and *τ* _*z*_).

When moths responded to pitch conditions, the time between the DLM spike and the first DVM spike increased in duration when pitching up (Fig. 3). In absolute time, mean duration between spikes across same-side DLM-DVM pairs was significantly different between pitch up and pitch down conditions (pitch up 24.5*±*0.8ms, pitch down 17.6*±*0.7ms) (Fig. 3A-B,E). Interestingly, wingbeat frequency was much lower in pitch up than pitch down conditions in all moths, with mean wingbeat frequency of 18.6*±*0.4Hz in pitch up and 22.9*±*0.5Hz in pitch down (Fig. 3F). This could indicate that moths lower their wingbeat frequency when executing pitch up maneuvers. Because there is a change in wingbeat period between pitch up and pitch down conditions of *≈*10.1ms which could account for the observed difference in *t*_*DVM*_ *−t*_*DLM*_ in absolute time, we normalized *t*_*DVM*_ *−t*_*DLM*_ to the length of each wingstroke (Fig. 3C-D,G). Mean phase between same-side DLM-DVM pairs was 45*±*1% of the wingstroke in pitch up conditions and 40*±*1% in pitch down conditions, with the majority of individuals showing this trend (Fig. 3G).

**Fig. 3.**
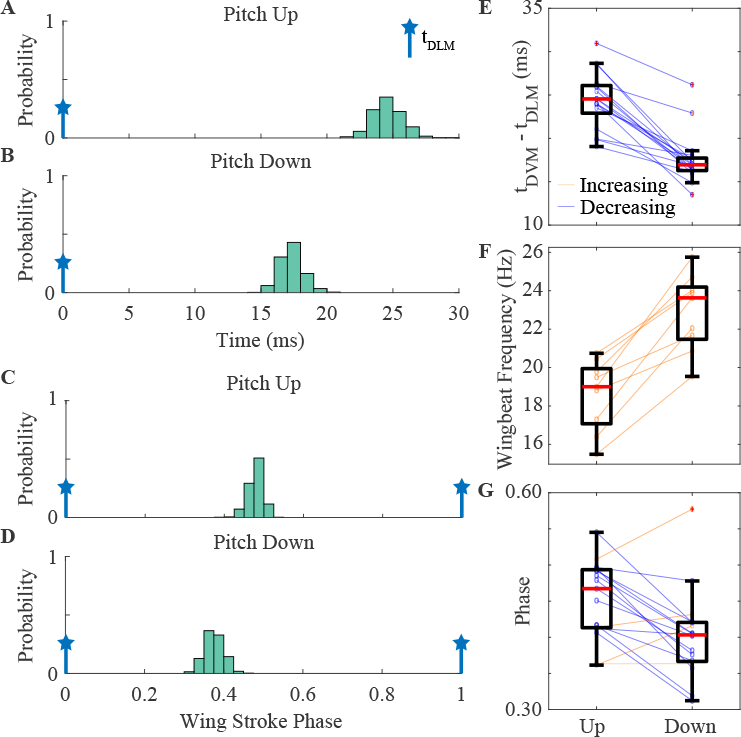
Timing between the DVM and DLM muscles increases when moths pitch up. (**A-B**) Normalized histograms from an example moth of the first DVM spike timing in each wing stroke (*t*_*DVM*_) with respect to the DLM timing (*t*_*DLM*_ = 0) for the pitch up and pitch down conditions. (**C-D**) Normalized histograms from the same example moth of the phase of the first DVM spike in each wing stroke with respect to the DLM timing. (**E**) Absolute timing difference between *t*_*DVM*_ and *t*_*DLM*_ for 17 same-side DLM-DVM pairs (*N* = 9 moths for 19 same-side pairs, with one failed DVM recording eliminating a pair). Means are statistically different, *p <* 10^*−*6^ for paired t-test. (**F**) Wingbeat frequency for all 9 moths for the pitch up and pitch down conditions (means statistically different, *p <* 10^*−*3^ for paired t-test). (**G**) Phase of the first DVM spike in each wing stroke with respect to the DLM timing for 17 same-side DLM-DVM pairs (means statistically different, *p* = 0.003 for paired t-test). Boxplots report the mean across individuals (red line), the 25th and 75th percentiles (box region), and the total range (whiskers) excluding outliers (red crosses).

Together these results demonstrate a correlation between DLM and DVM timing and pitch turns: In pitch up, the time between DLM and DVM spikes is greater, and in pitch down the time is reduced. While this data does not implicate any specific mechanisms or causal links between this timing shift and pitch turning, the high consistency of the observed timing shift suggests a functional role played by the power muscles in controlling pitch turns.

### Altered DLM timing produces consistent changes to within-wingstroke pitch torque dynamics

Tuned, brief electrical stimulation can elicit MAPs in both of the main downstroke muscles of *Manduca sexta* without cross-stimulation of other muscles (Fig. 2B), including in tethered flapping when all flight muscles are highly active (Fig. 2C-D). Beyond simply inducing spikes, this method of electrical stimulation can perform a “motor overwrite”, fully preventing natural spikes from occurring in their normal timing, if the natural spikes are suppressed by the refractory period of the motor unit

Causally manipulating DLM timing in single wingstrokes produced consistent and observable effects on within-wingstroke pitch torque (Fig. 4). When pitch torque traces are binned into groups by stimulation phase, pitch torque deviates from pre-stimulus wingstrokes (Fig. 4A-B). Typical wingstrokes have a 3- or 4-cycle oscillation in pitch torque which matches robotic flapping models of *Manduca* (Cheng et al., 2011), but evoked DLM spikes shifted the phase of these torque oscillations. Note that for all individuals, immediately after stimulation pitch torque initially pushes positive (head tilting downwards, as defined in Fig. 1A).

**Fig. 4.**
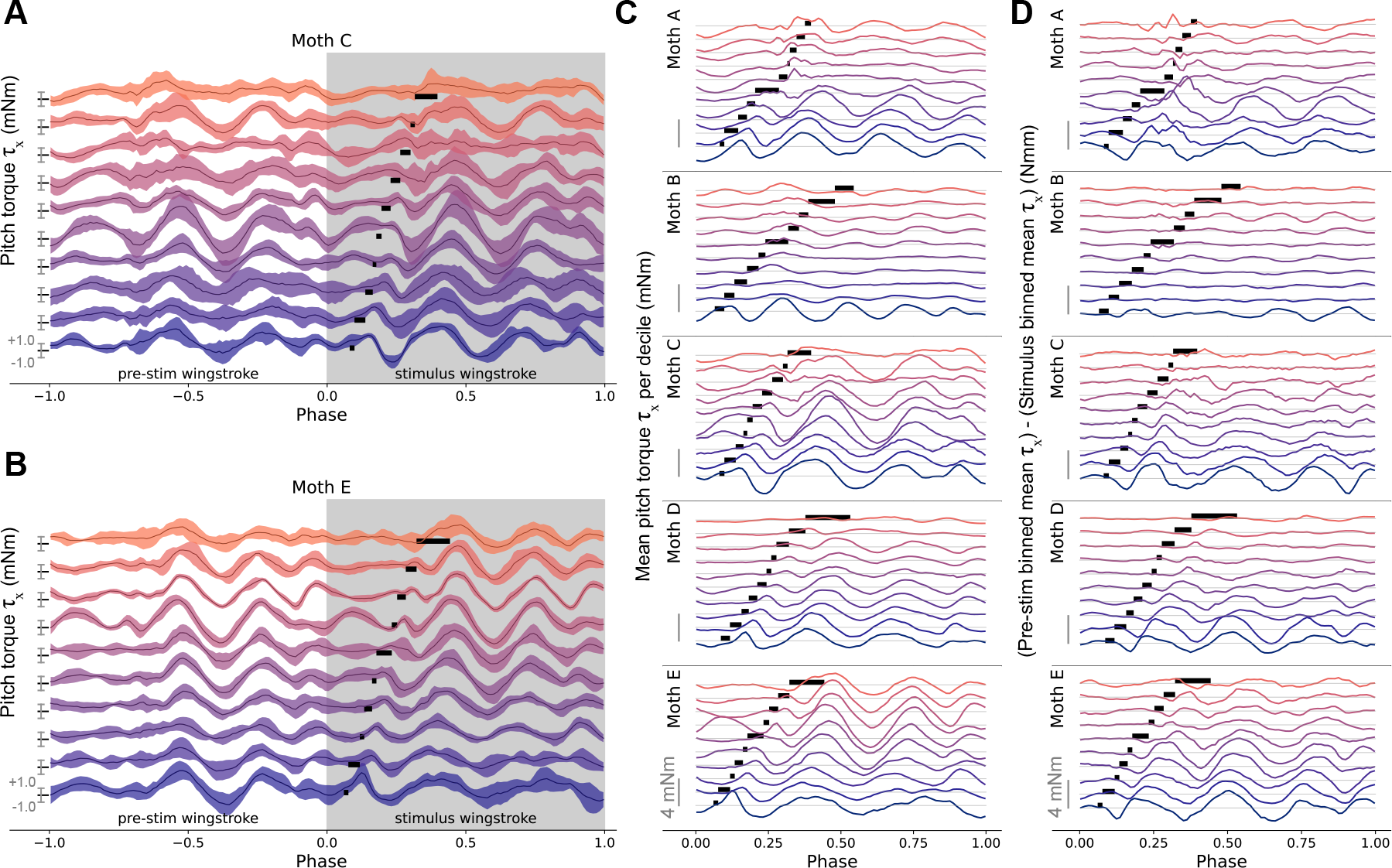
Qualitative features of stimulation visible in mean pitch torque waveforms binned by decile of stimulation phase. (**A**,**B**) Mean *±*1 standard deviation of time-varying pitch torque in the wingstrokes before, during, and after stimulation for two example individuals, binned by decile of stimulation phase. Each binned mean*±*S.D. trace contains the same number of pitch torque traces. Phase range during which stimulus was applied for each binned group is denoted by bar in stimulus wingstroke. Axes on left of each trace indicate where pitch torque (in mNm) is zero, +1, and *−*1 for each group. (**C**) Plots following(A) and (B) for all individuals, but showing only mean pitch torque for only the stimulation wingstroke. Light grey line indicates zero mNm for each binned group.(**D**) Difference in mean pitch torque during stimulated wingstroke compared to pre-stimulation wingstroke. At each phase the difference 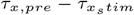 is taken, so this plot indicates the deviation from undisturbed, pre-stimulation pitch torque profile. Scale bars indicate 4mNm for each individual, light grey line indicates zero mNm for each binned group.

Stimulated wingstrokes show peak changes of 4mNm or more compared to the immediately preceding wingstroke (Fig. 4D). These changes lessen over the course of the wingstroke. Changes in torque were notably larger when the DLM was activated in early-wingstroke phases of less than 0.25. As a built-in control for the realism of evoked MAPs, for all individuals these changes due to stimulation progressively lessened as evoked DLM timing approached naturalistic timing around 40-50% of wingstroke phase.

Changing the relative timing between the activation of down-stroke and upstroke flight power muscles affects both mean pitch torque (Fig. 4C) and the deviation of mean pitch torque from normal dynamics (Fig. 4D). Altered DLM timing demonstrated consistent changes to within-wingstroke pitch torque dynamics in the form of a large oscillation in pitch torque following DLM spike phase, with the amplitude of this oscillation increasing the farther the DLM was evoked to spike relative to naturalistic timing. These features of causal manipulation of DLM timing, however, are purely qualitative. Surrounded by natural variation in pitch torque and the underlying motor program, features of pitch torque directly tied to evoked DLM timing are difficult to access and quantify without some form of directed feature extraction.

### CCA features demonstrate power muscles have mechanically relevant influence on pitch torque

To determine whether the power muscles have a controllable influence on pitch, and how much variation in pitch torque is causally attributable to power muscle timing, requires quantifying the specific features of pitch torque associated with DLM phase. CCA was applied to extract these features of pitch torque which maximally covaried with evoked changes in DLM phase (Fig. 5). CCA features pick out the same observed trend in Fig. 4, with the main cyclic oscillations of pitch torque shifting earlier in phase as the DLM was induced to spike earlier.

**Fig. 5.**
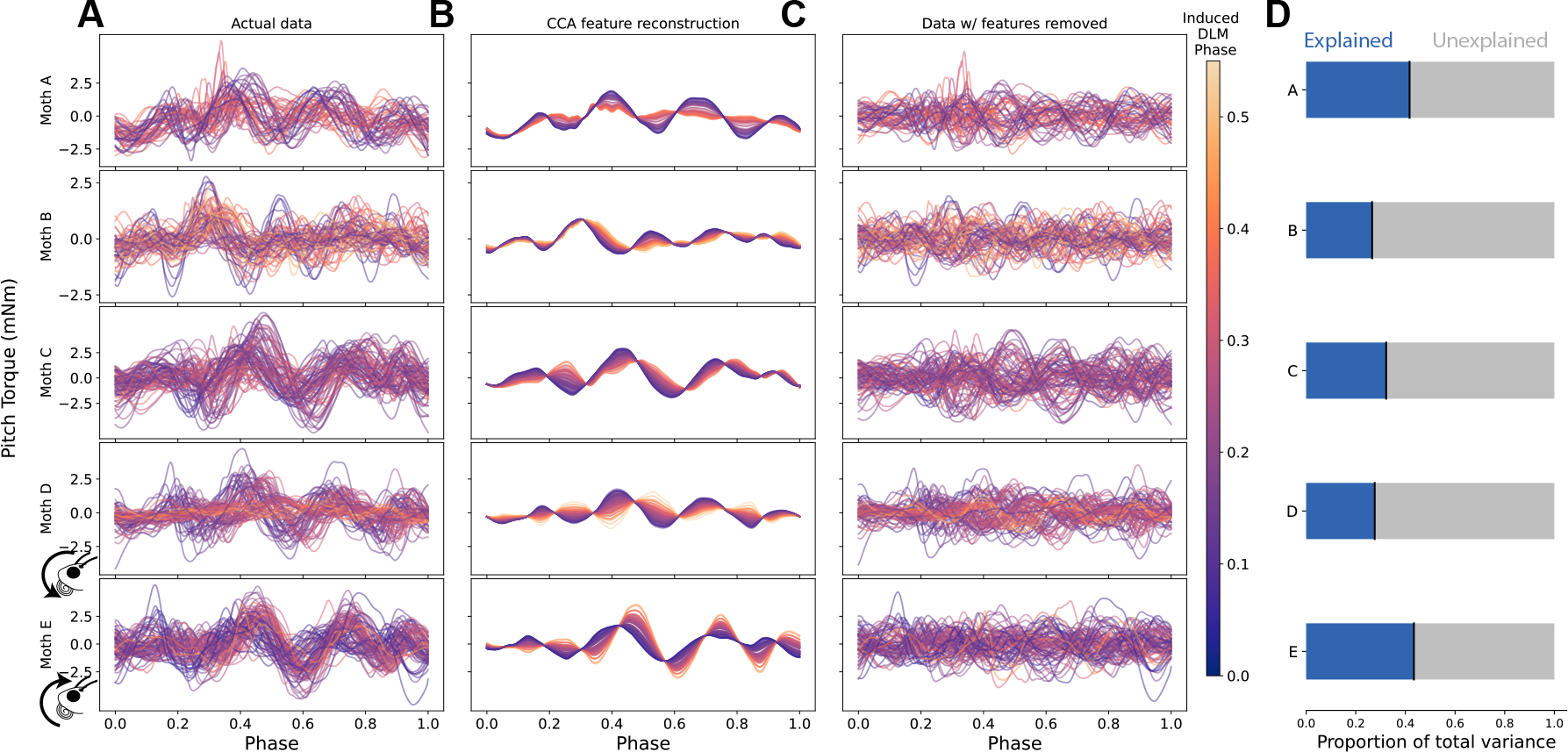
Features of pitch torque which covary with evoked DLM timing show consistent phase shift in pitch torque. (**A**) CCA feature reconstructions of pitch torque in stimulation wingstroke, colored by evoked DLM phase. Reconstructions generated using Eq. (1) followed by removal of z-scoring. (**B**) Actual data of pitch torque in stimulation wingstroke, colored by evoked DLM phase on the same scale as (A-D). (**C**) Raw data with CCA feature reconstructions subtracted, leaving only variance in pitch torque signal unexplained by CCA features. (**D**) Proportion of total variance explained by CCA feature reconstructions (blue, left) vs. unexplained (grey, right)

If DLM phase has a causal, controllable effect on body pitch, the angular impulse (change in angular momentum) in the stimulation wingstroke should vary significantly with DLM phase. The angular impulse in pitch imparted by reconstructed CCA features (Eq. (2)) varied linearly with DLM phase (Fig. 6A), with a statistically significant positive slope across individuals (linear mixed effects model with individual moth as a random intercept effect, *p <* 10^*−*3^). A significant slope is unsurprising, as CCA inherently finds features which maximally correlate with evoked DLM timing. CCA, however, is agnostic to the directionality of this relationship, so a positive slope indicates the maximally correlated relationship between these two variables is positive. Such a positive relationship indicates more negative angular momentum (more nose-up pitch) is linearly associated with a greater interval between the DLM and DVM, matching observations of the correlational experiment where greater *t*_*DVM*_ *−t*_*DLM*_ was associated with pitching up (Fig. 3E).

**Fig. 6.**
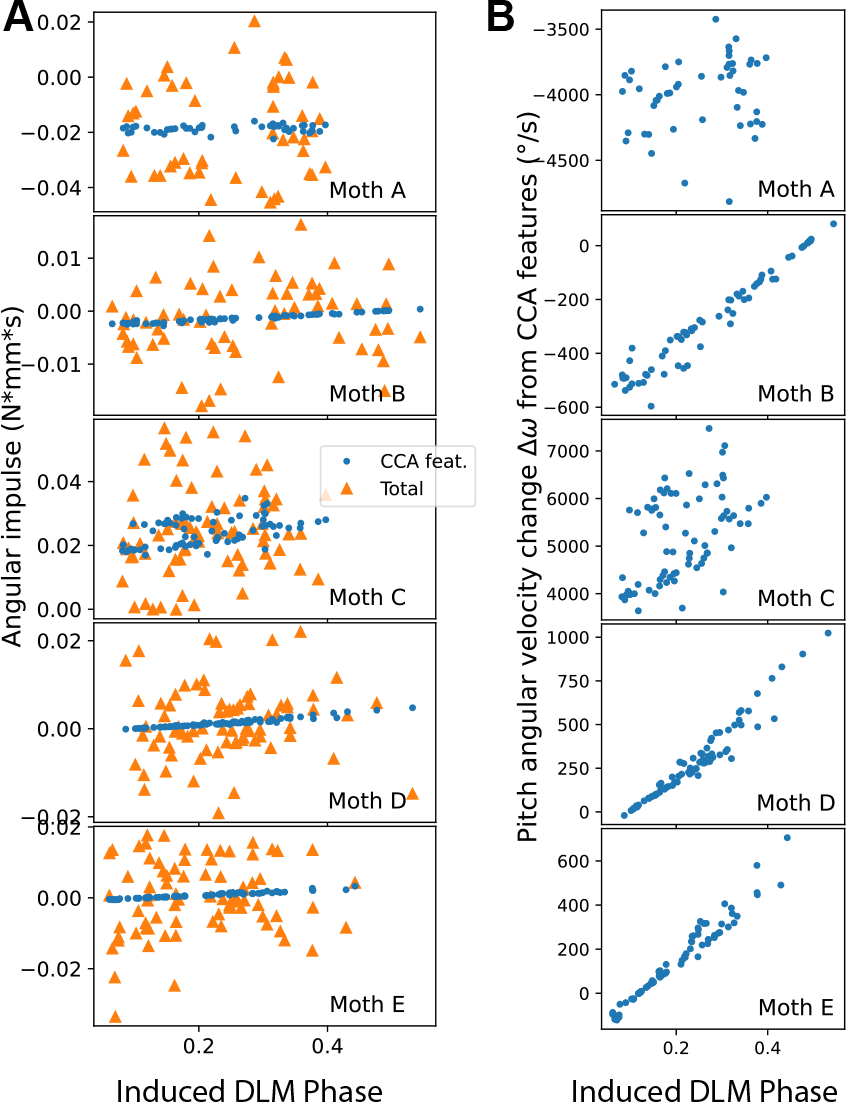
CCA features produce mechanically relevant levels of angular impulse in pitch. (**A**) Angular impulse from CCA feature reconstructions of pitch torque associated with DLM phase (blue, circles, Fig. 5B) and overall pitch torque (orange, triangles, Fig. 5A) for each individual moth. (**B**) Pitch angular velocity change due to angular impulse from CCA feature reconstructions, in degrees per second.

To assess the behavioral relevance of these pitch torques, we divide angular impulse by pitch moment of inertia *I*_*yy*_, giving the effective change in pitch angular velocity Δ*ω* over a wingstroke (Eq. (3)). Change in pitch angular velocity over a wingstroke due to reconstructed features increased by at least 400 °*/s* between early and late DLM phase in every individual (Fig. 6B). For a typical 50 millisecond wingstroke, 400 °*/s* corresponds to a change of at least 20° in body pitch angle over a single wingstroke, a relevant amount of change from a flight control perspective. Pitch angular velocity of 400 °*/s* is also well in line with free-flight pitching maneuvers pitch angular velocities observed to vary between -500 to +500 °*/s* in *Manduca* (Cheng et al., 2011).

While it is clear that the features of pitch torque driven by the DLM demonstrate a causal and mechanically relevant influence on body pitch, there is a notable amount of unexplained variance of pitch torque in the stimulation wingstroke (Fig. 5D). On average across individuals, 64.5*±*6.7% of the variance of pitch torque is not explained by a linear feature of evoked DLM phase. Similarly, while CCA features produced mechanically relevant angular impulse, the magnitude of impulse due to these features is far smaller than overall pitch angular impulse (Fig. 6A). CCA does not seek features which maximize explained variance or angular impulse, instead only seeking features which maximally covary with the change in timing. Leaving large amounts of unexplained variance, then, is expected. However, this indicates that the degree to which muscles other than the DLM influence pitch torque is high, and the DLM does not influence pitch in isolation.

## DISCUSSION

### Power muscles contribute to pitch control in hawkmoth flight

The two experiments of this paper present correlational and causal evidence which supports the hypothesis that bilaterally symmetric changes in DLM spike timing alter body pitch. Greater time between the DLMs and DVMs leads to upwards pitch, and less time leads to downwards pitch. In the first experiment, pitch turns were visually induced and verified to produce separated responses in mean-centered torques only about the pitch axis (Fig. 1), and decreases in the phase between the DLM and DVM were observed in pitch down turns compared to pitch up (Fig. 3). This correlational finding was corroborated through causal manipulation of DLM-DVM timing (Fig. 2), where features of pitch torque that covaried with evoked DLM timing shifted towards downwards mean pitch as the time between the DLM and DVM decreased (Fig. 5). Bilaterallly asymmetric power muscle timing has been linked to control of yaw turns in flies and hawkmoths (Sponberg and Daniel, 2012; Tu and Daniel, 2004; Lehmann et al., 2013), but to date no studies have shown symmetric power muscle timing can causally affect pitch. This is a particularly relevant finding as pitch is part of an are inherently unstable dynamic mode, and requires active control on relatively short timescales on the order of *≈*45*ms* for *Manduca* (Ristroph et al., 2013), similar to the duration of a typical wingbeat of *≈*50*ms*.

The ability for the indirect flight muscles to control pitch does not preclude known contributions from direct steering muscles in species including *Drosophila* (Whitehead et al., 2015; 2022) and *Manduca* (Ando and Kanzaki, 2004). This is consistent with induced changes in DLM timing relative to DVMs producing consistent pitch torque features, but the variance explained by these features being relatively low (Fig. 5D) and the angular impulse of these features, while relevant, presenting as small compared to overall impulse (Fig. 6).

A smaller influence of the indirect power muscles on pitch, however, is still behaviorally relevant. Despite often being much smaller than total pitch impulse, impulse from CCA features was large enough in all individuals to cause 400°*/s* or more change in pitch angular velocity over single wingstrokes (Fig. 6). Both the amount of pitch imparted by CCA features and the ranges of DLM timing required to do so are highly plausible in free flight. The observed 400 °*/s* pitch velocity change closely matches the *±*500°*/s* pitch velocity observed in *Manduca* free flight (Cheng et al., 2011), and in natural, unmanipulated wingstrokes the DLM spikes between 20% to 55% of the wingstroke phase (Putney et al., 2019). So even though the most pronounced effects of causally manipulating DLM phase on pitch torque occurred in earlier phases near 20-30% (Fig. 4D), these phases are consistent with the natural timing range of the DLM. Thus, while the overall effects of the DLM may only explain *≈*40% of the variation in pitch torque, this is a relevant amount of variation when it comes to controlling the unstable mode of pitch as the observed effects on pitch from the DLM are reasonably expected to occur in natural free flight.

### Possible mechanisms for power muscles to modulate pitch

For a flapping wing animal such as an insect, there are four possible ways to produce and modulate body pitch torque: 1) Move the location of the body center of mass (COM) relative to the wing center of pressure (COP) to change the moment arm on which aerodynamic forces are applied (e.g. abdominal flexion).2) Modify the location of the COP in relation to the COM by altering wing kinematics (e.g. changing wing elevation angle). 3) Alter wing kinematics to produce inertial moments on the COM. 4) Change the within-wingstroke aerodynamic forces produced by the wings (e.g. changing the duration of the downstroke and upstroke to exploit moment asymmetries between the two phases of the wingstroke). Of these four mechanisms, the indirect power muscles are only able to alter wing kinematics, meaning the most likely mechanisms to modulate pitch are through movement of the COP, changing time-varying wing forces, and modifying inertial moments.

For indirect power muscles, the most plausible kinematic mechanisms to exert pitch control are either by changing time-varying wing forces or moving the COP, both of which have been observed in hawkmoths. In freely flying hawkmoths feathering angle, or the average rotation of the wing about its base-to-tip axis, correlates directly with pitching movements (Cheng et al., 2011). As confirmed by a model robotic flapping wing, varying feathering angle has a clear effect of altering time-varying wing forces by altering the wing angle of attack. Increase mean feathering angle, and the angle of attack is increased during downstroke and decreased during upstroke, leading to a net nose-up pitching moment (Cheng et al., 2011). There is little direct path, however, for power muscles to alter wing feathering angle, a kinematic quantity thought to be primarily controlled by direct steering muscles such as the 3^rd^ axillary muscle in hawkmoths (Ando and Kanzaki, 2004; Rheuben and Kammer, 1987).

Moving the COP relative to the COM is a more likely method for power muscles to modulate pitch. Tethered hawkmoths use this method to induce pitch turns in response to looming stimuli, altering the mean wing elevation angle to shift the angular extents the wing achieves in the wing stroke plane and thus shift the COP (Ando and Kanzaki, 2016). The relative timing between the DLM and DVM is known to alter the wingstroke amplitude and elevation angle along the stroke plane from wingstroke to wingstroke in hawkmoths (Wang et al., 2008), and provides a plausible change to the COP location which can induce a pitching moment.

There is precedent for pitch control via bulk wingstroke elevation angle adjustment in more flying insects than just hawkmoths. In flies changing mean wing elevation angle is essential for pitch control (Dickinson, 1999; Whitehead et al., 2015; 2022). Previous work in flies has also connected wing rotation angle to pitch control (Whitehead et al., 2015; 2022), as well as implicated additional mechanisms not directly observed in hawkmoths such as tilting of the wingstroke plane (Zanker, 1988) and wingstroke frequency (Dickinson, 1999). In lepidopterans, while indirect power muscles can certainly alter wingstroke frequency, and are well-understood to generate wing motion on the major stroke plane (Kammer, 1985), there is little direct evidence for power muscles demonstrating a control potential on wing rotation angle or stroke plane tilt. For this reason, of the possible kinematic mechanisms listed we find it most likely that power muscles exert pitch control authority through wing elevation angle and possibly wingstroke frequency.

Regardless of the mechanism, any pitch control potential from power muscles must happen in concert with the control potential of direct steering muscles. In flies the 1^st^ and 2^nd^ basalar muscles likely contribute to the control of wing elevation and wingstroke amplitude on short timescales (Whitehead et al., 2015; 2022; Lindsay et al., 2017; Heide and Götz, 1996; Tu and Dickinson, 1996; Lehmann and Götz, 1996; Balint and Dickinson, 2001). While in synchronous hawkmoths there is a strong case for indirect power muscle control potential of yaw turning maneuvers, (**?**Sponberg et al., 2015a; Putney et al., 2019), this does not preclude the influence of the rest of the flight musculature. Our results demonstrate that the indirect power muscles have some control potential on body pitch, but this control potential only occurs in the context of the rest of the motor program and flight musculature.

### Precisely changing the timing of one power muscle doesn’t produce a precise turn

Of important note is the relatively high amount of unexplained variance left after removal of CCA features (Fig. 5C-D, 64.5*±*6.7% across individuals). While CCA does not extract features which maximize variance in a dataset, only seeking features which maximize covariance between pitch torque and evoked DLM timing, the large amount of pitch torque signal unexplained by evoked DLM timing leaves an interesting implication. Despite the DLM being the largest muscle in *Manduca sexta*, large changes to DLM timing did not lead to large changes in tethered flight dynamics. Stated differently, the timing of one power muscle alone doesn’t produce a turn.

Our results put a precise understanding on where, between the two extremes of functionally overlapping and functionally separated, the hawkmoth flight power musculature falls. By using causal electrical stimulation in concert with dual dimensionality reduction, we directly measured the degree to which a specific behavioral output, pitch torque, is driven by the control potential of a single pair of muscles, the DLMs. The DLMs do exert a measurable control potential on pitch torque sufficient to produce pitch turns (Fig. 4D, Fig. 6B). But the features of pitch torque associated with the timing of the DLMs fail to explain most of the variation in pitch torque and poorly describe pitch angular impulse in wingstrokes (Fig. 5D, Fig. 6A). This result shows that in controlling body pitch, at least some subset of the flight musculature *must* be coordinated to produce a pitch turn. Even if all the flight muscles have fully separate and non-overlapping kinematic functions, to produce the results observed these muscles must have some functional overlap in pitch torque.

This fits well with prior results in hawkmoths and other flying insects, where individual behavioral functions of individual muscles can often cotnribute to specific actions in isolation, yet measurable functional overlap and interactions between muscles seem to require many coordinated muscles to describe behavior. In flies the 1^st^ and 2^nd^ basalar muscles are frequently correlated to stroke amplitude (Whitehead et al., 2015; 2022; Lindsay et al., 2017; Heide and Götz, 1996; Lehmann and Götz, 1996; Balint and Dickinson, 2001). But even these two muscles are known to have interacting kinematic effects (Tu and Dickinson, 1994; 1996; Dickinson and Tu, 1997), and the activity of at least 8 of 12 steering muscles is required to describe more than 30% of the variance of just wing stroke amplitude (Lindsay et al., 2017). In hawkmoths, only activity from any 4 or more muscles is needed to decode the type of turn being performed with more than 90% accuracy (Putney et al., 2021a). Similarly, redundant global information in spike timing has been observed across the entire flight motor program (Putney et al., 2019), suggesting functional coordination such that individual muscles are controlled in the context of the rest of the motor program. Hawkmoths do indicate some aspects of a functionally separated flight musculature, however. Compared to functionally coordinated systems where muscle synergies, typically linear combinations of motor commands across multiple muscles, can describe the majority (often >90%) of output variation (Ting, 2007), yaw turning behaviors in hawkmoths were better described by the independent activity of muscles than as a synergy (Sponberg et al., 2015a).

Our results also provide a causal test of the temporal precision that has been estimated from information theoretic analysis of the hawkmoth flight motor program (Putney et al., 2019; 2021b). Changes to DLM timing were applied during tethered but unconditioned behavior; moths were not cued or tuned to any particular behavior or maneuver other than wing flapping, and thus DLM timings were applied across a distribution of locomotor coordination states. Precise changes in DLM timing did lead to precise changes in the CCA feature space (Fig. 6B), but these small changes were typically only a portion of the variation in the rest of the motor program (Fig. 6A). Thus, precision is relative to coordination. While the spike timing of a muscle may describe the yaw torque of a hawkmoth to a sub-millisecond scale (Putney et al., 2021b), this is only in the context of the rest of the motor program also aiming to produce that yaw torque. When muscles functionally overlap to control an outcome like pitch or yaw torque, then the specific timing of one muscle is context-dependent, with an impact which can be cancelled out by other muscles.

Overall, then, our results underline the importance of coordination in flight musculature to produce controlled flight. While kinematic or behavioral outcomes can be causally attributed to the spike timing of individual muscles, there is enough functional overlap between muscles in the flight motor program to make the effects of muscle spike timing context-dependent. Even the largest and most powerful muscles in a hawkmoth have only so much explanatory power when manipulated outside of the coordinated activity of all other muscles.

## Data availability

Visually-induced hard turns data is available on Dryad (Putney et al., 2023).

## Supplementary

No supplementary text included.

